# Phosphoproteomic Profiling Reveals Overlapping and Distinct Signaling Pathways in *Dictyostelium discoideum* in Response to two different chemorepellents

**DOI:** 10.1101/2025.10.21.683751

**Authors:** Salman Zahir Uddin, Ramesh Rijal, Richard H. Gomer

**Author notes:** Address correspondence to: Richard H. Gomer, Department of Biology, Texas A&M University, ILSB 301 Old Main Drive, College Station, TX 77843-3474 USA.

## Abstract

Chemorepulsion causes cells to move away from the source of a signal, but the underlying mechanisms for eukaryotic cells are poorly understood. We performed proteomics and phosphoproteomics to elucidate how *Dictyostelium discoideum* responds to its two endogenous chemorepellent signals, the protein AprA and inorganic polyphosphate (polyP). At 60 minutes, AprA increased levels of 211 proteins and reduced levels of 57 proteins while polyP increased levels of 152 proteins and reduced levels of 168 proteins. Surprisingly, many of the AprA- and polyP-regulated proteins are associated with RNA metabolism and ribosomes. AprA and polyP both upregulated 19 proteins, one protein was downregulated by both, and one was upregulated by AprA and downregulated by polyP. AprA increased phosphorylation of 12 proteins and decreased phosphorylation of 60 proteins. PolyP increased phosphorylation of 7 proteins and decreased phosphorylation of 18 proteins. As expected, the two chemorepellents affected phosphorylation of signal transduction/ motility proteins, but unexpectedly affected phosphorylation of RNA-associated proteins. Both AprA and polyP decreased phosphorylation of six proteins including the Ras-interacting protein RipA and guanine nucleotide exchange factors (GEFs) such as the RacGEF GxcT. Mutants lacking RipA or GxcT were unresponsive to both AprA and polyP chemorepulsion. Together, this work supports the idea that rather than activating the same chemorepulsion mechanism, AprA and polyP activate only partially overlapping chemorepulsion mechanisms, and identifies two new components that are used by both chemorepellents.

## Introduction

Directed cell migration is a fundamental biological process essential for morphogenesis, tissue repair, and immune surveillance (SenGupta et al., 2021; Shellard & Mayor, 2020). While the molecular mechanisms of chemoattraction, the process by which cells migrate toward stimuli have been extensively studied, the opposite behavior, chemorepulsion, remains comparatively underexplored (Fu & Feng, 2024; SenGupta et al., 2021). Chemorepulsion enables cells to migrate away from harmful or crowded environments and plays vital roles in immune regulation, inflammation resolution, and spatial patterning during development (Jin, 2023; Kirolos et al., 2021).

The social amoeba *D. discoideum* is a well-established eukaryotic model organism for investigating chemotaxis due to its genetic tractability and the conservation of many signaling pathways with higher eukaryotes (Bozzaro, 2019; Pears, 2021). In this system, two endogenous chemorepellents have been identified: AprA, a ∼60 kDa secreted autocrine protein, and inorganic polyphosphate (polyP), a polymer of phosphate (El-Sobky et al., 2025; Kirolos & Gomer, 2022; Phillips & Gomer, 2012; Rijal et al., 2019; Tang et al., 2018). The two repellents can function independently, but both cause cells at the edge of a colony to move away from the colony, presumably to find sources of food. In *D. discoideum* and other systems, a localized activation of Ras at one sector of the cell membrane activates pseudopod formation and movement in the direction of the pseudopod (Artemenko et al., 2014; Huang et al., 2013; Matsuoka et al., 2024). A gradient of AprA induces repulsion by inhibiting Ras activation and pseudopod formation at the side of the cell closest to the source of AprA, causing the cell to move in any direction except toward the source of AprA (Kirolos & Gomer, 2022; Rijal et al., 2019). In contrast, a gradient of polyP induces chemorepulsion by activating Ras at multiple cortical sites and increasing pseudopod formation, particularly at the side of the cell furthest from the polyP source (El-Sobky et al., 2025).

Although both AprA and polyP can trigger repulsion, their signaling mechanisms only partially overlap. AprA signals through the G protein-coupled receptor (GPCR) GrlH, activating the heterotrimeric G-protein subunits Gα8 and Gβ, which in turn signal through PakD, CnrN, TORC2, Erk1, and the Ras proteins RasC and RasG (Bakthavatsalam et al., 2009; Phillips & Gomer, 2012; Rijal et al., 2019; Tang et al., 2018). PolyP signals through the GPCR GrlD, using different G-protein subunits, and uses PI3 kinases, phospholipase C, and effectors such as WasA and NapA to control actin dynamics (El-Sobky et al., 2025; Suess et al., 2019; Tang et al., 2021). While AprA requires both RasC and RasG for repulsion, polyP relies solely on RasC, underscoring their differences in small GTPase regulation despite sharing other core signaling elements (El-Sobky et al., 2025).

In addition to inducing repulsion, AprA and polyP both inhibit cell proliferation (Choe et al., 2009; Suess & Gomer, 2016). AprA-mediated proliferation inhibition involves RasC/RasG and Erk1 signaling pathway components also required for chemorepulsion, but requires some signal transduction pathway components not needed for repulsion to inhibit proliferation (Kirolos et al., 2021; Phillips & Gomer, 2010; Phillips et al., 2011). PolyP also uses some components needed for repulsion to also inhibit proliferation, and has at least one pathway component not needed for repulsion that is used to inhibit proliferation (El-Sobky et al., 2025; Suess et al., 2019; Tang et al., 2021).

Many extracellular signals affect phosphorylation of specific proteins (Jin, 2023; Ridley et al., 2003; Swaney et al., 2010). Our understanding of chemorepulsion pathway components in *D. discoideum* only used known mutants. To gain a broader perspective, in this study we used proteomics and phosphoproteomics to elucidate the effects of AprA and polyP on cells, and then examined the role of two selected proteins whose phosphorylation appeared to be regulated by AprA or polyP. We find that AprA and polyP affect levels and phosphorylation of partially overlapping pathway components centered on small GTPase signaling, and show that two proteins whose phosphorylation is reduced by AprA and polyP are necessary for the effects of both chemorepellents on repulsion but not proliferation.

## Materials and Methods

### Cell strains and culture

*D. discoideum* strain Ax2 (Dictybase identifier DBS0237699) (Watts & Ashworth, 1970) was obtained from the Dictyostelium stock center (Fey et al., 2013). The strains *ripĀ* (DBS0236900) (Rosel et al., 2012), and *gxcT^−^* (DBS0350268) (Wang et al., 2013) were gifts from Dr. Alan Kimmel (Laboratory of Cellular and Molecular Biology, NCI, NIH, Bethesda, MD, USA) and Dr. Miho Iijima (Department of Cell Biology, Johns Hopkins University School of Medicine, Baltimore, MD, USA) respectively. Cells were cloned on SM/5 agar with DB *Escherichia coli* (DBS0350636) and then grown in HL5 medium (#HLG0101, Formedium, Hunstanton, UK) containing 100 µg/ml dihydrostreptomycin (#S-150-50, GoldBio, St. Louis, MO, USA) and 100 µg/ml ampicillin (#A-301-25, GoldBio) as previously described (Fey et al., 2007; Paschke et al., 2018). For HL5 growth, cells were either in submerged culture on 10 cm tissue culture dishes (#10861-680, Avantor, Randor, PA, USA) or in suspension at 22 °C with continuous shaking at 180 rpm (Fey et al., 2007; Watts & Ashworth, 1970). Mutants were grown under constant selection with 5 µg/ml blasticidin (#B-800-25, GoldBio) in both submerged and shaking culture.

### Stimulation with AprA and polyP

Ax4 cells were grown in 50 ml of HL5 medium at 22 °C with shaking at 180 rpm until mid-log phase (3 × 10⁶ cells/mL) (Fey et al., 2007; Watts & Ashworth, 1970). Cells were collected by centrifugation and washed twice with 10 ml of HL5 at 380 × g for 5 minutes at 4 °C. Cells were then resuspended in HL5 to 1 × 10⁶ cells/ml. For AprA treatment, freshly purified recombinant AprA (produced using His-tag affinity chromatography as described in (Bakthavatsalam et al., 2009) was added to a final concentration of 300 ng/ml. For polyphosphate (polyP) stimulation, sodium polyphosphate (average chain length ∼46; #S0169, Spectrum Chemical, New Brunswick, NJ) was freshly prepared as a 70 mg/ml stock in water and added to a final concentration of 210 μg/ml (Suess & Gomer, 2016). Before adding AprA or polyP (0 minutes), and at 60 minutes, 3.5 ml aliquots were transferred into pre-chilled 15 ml conical tubes (Corning, Corning, NY, USA), collected by centrifugation at 2000 × g for 5 minutes at 4 °C, and placed on ice. Pellets were resuspended in 150 µl of ice-cold RIPA buffer (#PI89901, Thermo Fisher Scientific, Waltham, MA, USA) supplemented with Halt™ phosphatase inhibitor cocktail (#78420, Thermo Fisher Scientific, Waltham, MA, USA) and complete protease inhibitor cocktail (#11836170001, Roche Diagnostics, Basel, Switzerland). Lysates were incubated on ice for 30 minutes with occasional mixing to enhance extraction efficiency. Following centrifugation at 18,000 × g for 10 minutes at 4 °C, 140 µl of the clarified supernatants were transferred to pre-chilled Eppendorf tubes. 10 μl aliquots were set aside for protein concentration determination and the remaining supernatant was flash-frozen in liquid nitrogen and stored at −140 °C (Fey et al., 2007; Rijal & Gomer, 2024).

### Proteomic and phosphoproteomic sample preparation and analysis

Proteomics and phosphoproteomics were carried out at the University of Texas Southwestern (UTSW) Proteomics Core Facility (Dallas, TX, USA). Samples were shipped on dry ice to UTSW, where all sample processing, including protein quantification, reduction and alkylation, enzymatic digestion, phosphopeptide enrichment using titanium dioxide (TiO₂) beads, and peptide desalting, was performed following their standard protocols (Rappsilber et al., 2007; Shevchenko et al., 1996; Thingholm et al., 2006). LC-MS/MS analysis was performed on an Orbitrap Fusion mass spectrometer (Thermo Fisher Scientific, Waltham, MA, USA) using standard gradient methods. Raw data were processed and searched against the *D. discoideum* UniProt protein database using MaxQuant software (v1.6.17), applying a 1% false discovery rate at peptide and protein levels (Cox & Mann, 2008; Tyanova et al., 2016). Raw and processed proteomic data was uploaded to MassIVE at the University of California at San Diego Center for Computational Mass Spectrometry with accession number MSV000098894.

### Data normalization and analysis

Protein and phosphoprotein abundances were normalized within each replicate by dividing each individual abundance value by the total protein abundance of that replicate, thereby controlling for differences in sample loading and instrument response for both the total proteome and the phosphoproteome (Karpievitch et al., 2012). Log₂ fold changes were then calculated as the ratio of normalized abundance at 60 minutes relative to the normalized abundance at 0 minutes for the same protein or phosphoprotein. Proteins or phosphoproteins with zero abundance in all three replicates at a given time point were excluded from fold-change and statistical analyses; we identified two such proteins under polyP treatment and one under AprA treatment across both the time points, which are reported separately in the Results. Apart from these explicitly reported cases, all other proteins and phosphoproteins were included in the downstream analyses. Statistical significance was assessed using paired two-tailed Student’s t-tests on the normalized replicate values for each condition. Volcano plots were generated using Prism 10.4.0 (GraphPad Software, San Diego, CA, USA). Functional enrichment analysis of significant proteins and phosphoproteins was performed using ShinyGO v0.85 (Ge et al., 2020) with parameters set to an FDR cutoff of 0.05, a pathway size range of 5–1000 genes, and a maximum of 30-40 pathways displayed. Gene Ontology (GO) terms, and SMART domain annotation (Letunic et al., 2021) were included in the enrichment workflow.

### Protein annotation using AlphaFold and Foldseek

To assign putative functions to uncharacterized protein hits identified in our proteomic and phosphoproteomic analyses, we conducted structural homology searches using AlphaFold (Jumper et al., 2021; Varadi et al., 2022) and Foldseek (van Kempen et al., 2024). For both AprA- and polyP-treated samples, only proteins exhibiting greater than twofold changes in abundance or phosphorylation were selected for characterization. We also used Alphafold/Foldseek to characterize 7 proteins whose phosphorylation was affected by both AprA and polyP.

### Functional validation assays

Chemorepulsion assays were performed in Insall chambers as previously described (Muinonen-Martin et al., 2010; Phillips & Gomer, 2012; Rijal et al., 2019). HL5 medium was added to the central well, while either AprA or polyP diluted in HL5 was placed in the outer well, with an equal volume of HL5 media used as a control. Time-lapse images were collected every 15 seconds for 60 minutes using an inverted microscope. Migration parameters including forward migration index (FMI), speed, and persistence were quantified using the ibidi Chemotaxis and Migration Tool 2.0 (Stand-alone application; ibidi GmbH, Gräfelfing, Germany).

Proliferation inhibition assays were performed as described (Bakthavatsalam et al., 2009), with cells cultured at 5 × 10⁵ cells/ml in HL5 medium and treated with or without 300 ng/ml AprA or 700 µg/ml polyP. Viable cell counts were determined at 24 hours using a hemocytometer, and proliferation was calculated as a percentage of the untreated control for each replicate.

### Statistical analysis

Statistical analyses were performed using Prism 10.4.0 (GraphPad Software, Boston, MA). P < 0.05 was considered significant.

## Results

### AprA and polyP have mostly different effects on protein levels and phosphorylation

To investigate how AprA and polyP modulate signaling in *D. discoideum*, we first assessed effects on the proteome of proliferating cells at 60 minutes. We previously observed that it takes about 30 minutes for cells to respond to AprA or polyP, and that there is good repulsion at 60 minutes, so we chose this time point. Out of 5,383 proteins quantified (**Supplementary file S1**), AprA increased levels of 211 proteins, including 13 (**Supplementary file S1, tab 4**) with more than 2-fold increases and decreased levels of 57 proteins (**Figure 1A** and **Supplementary file S1, tab 3**). A previously uncharacterized protein (Q550S3) was upregulated by more than 2-fold at 60 minutes by AprA. Annotation by BLAST, AlphaFold, and FoldSeek suggested strong similarity to PhoPQ-activated pathogenicity-related proteins and to AprA-like isoforms **(Supplementary file S1, tab 4**). Another protein (MAP3 kinase 12 inhibitory protein; **Supplementary file S1, tab 5**), had zero abundance in all three replicates at 0 minute but was detectable at 60 minutes and was therefore excluded from fold-change and statistical calculations. At 60 minutes, polyP upregulated 152 proteins and downregulated 168 proteins (**Figure 1B** and **Supplementary file S2, tab 3**). Two proteins had zero abundance in all three replicates at 0 minute, were excluded from fold-change and statistical calculations but are listed in **Supplementary file S2**, **tab 4**. At 60 minutes, comparison of AprA- and polyP-treated proteomes revealed 19 proteins which were upregulated by both treatments, one (ubiquitin carboxyl-terminal hydrolase) was downregulated by both, and one (a non-specific serine/threonine protein kinase) was upregulated by AprA but downregulated by polyP (**Figure 2A; Supplementary file S7, tab 2**). For both AprA and polyP, many proteins did not exhibit significant changes, indicating a targeted rather than a global response.

**Figure 1.**
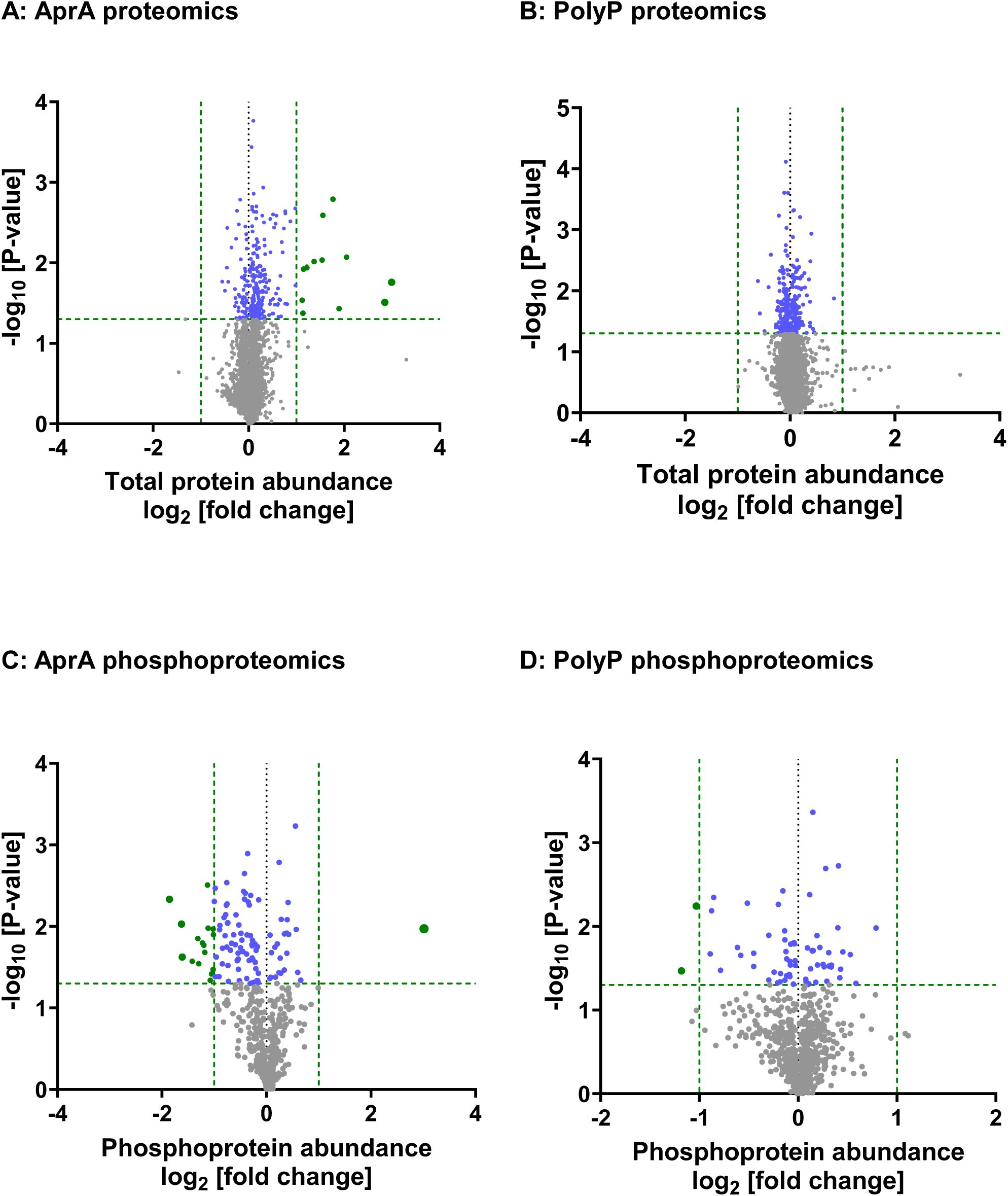
Comparative analysis of proteomic and phosphoproteomic changes induced by AprA and polyP. Volcano plots of protein abundance changes at 60-minutes post stimulation (x-axis: log₂ fold change; y-axis: −log₁₀ p). (**A**) Proteomic changes following AprA stimulation (n=5,383 proteins). (**B**) Proteomic changes following polyP stimulation (n=5,015 proteins). (**C**) Phosphoproteomic changes following AprA treatment (n=602 phosphoproteins). (**D**) Phosphoproteomic changes following polyP treatment (n=846 phosphoproteins). Green vertical dashed lines mark the significance threshold (log₂ fold change = ±1, equivalent to a ≥ 2-fold change in abundance), while the horizontal dashed line marks the statistical significance cutoff (p < 0.05). Green dots indicate proteins with |log₂ fold change| ≥ 1 and p < 0.05; blue dots indicate proteins with |log₂ fold change| < 1 and p < 0.05; grey dots indicate non-significant proteins.

**Figure 2.**
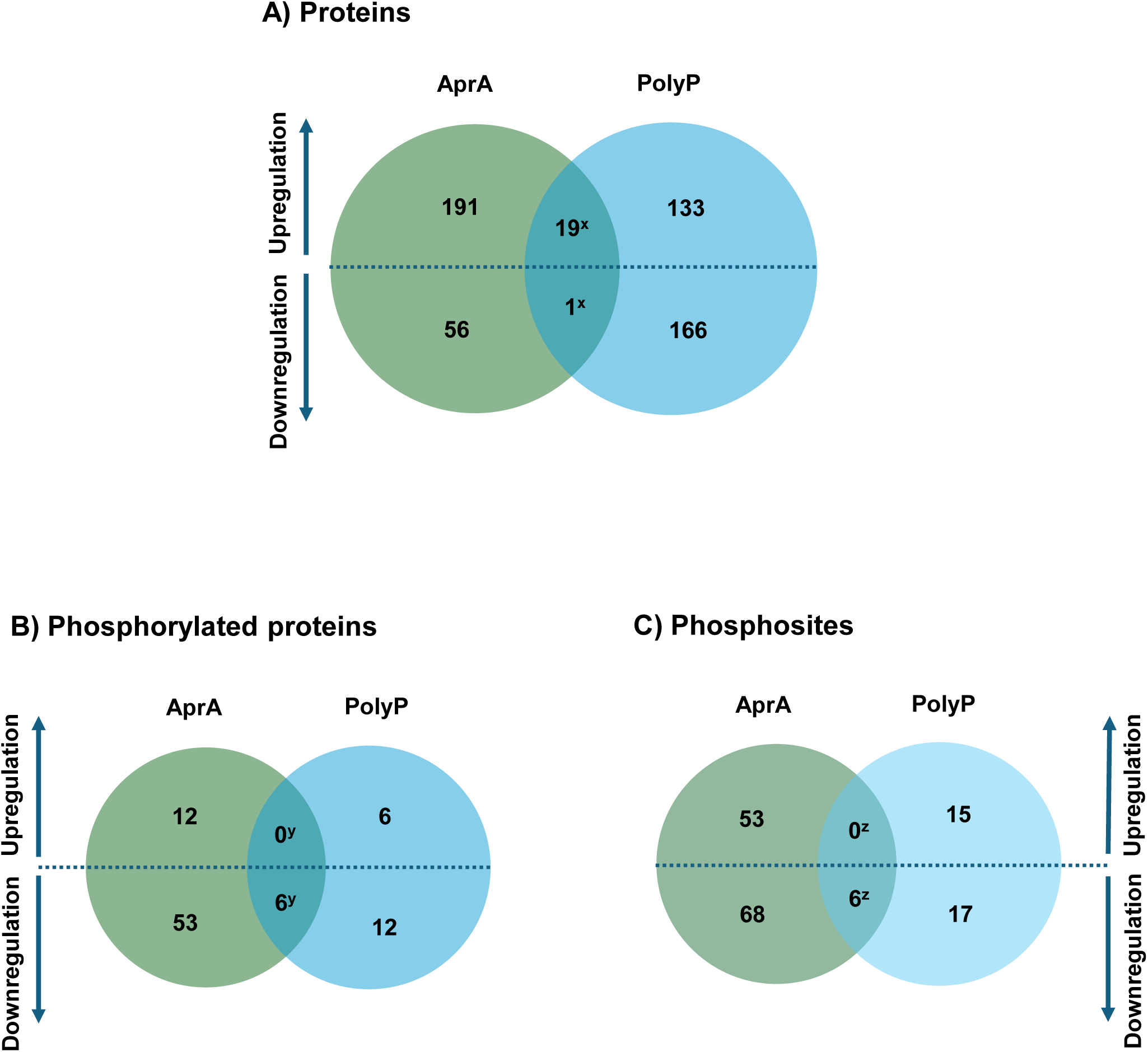
Proteins and phosphoproteins regulated by AprA and polyP. (**A**) Venn diagram of significantly regulated proteins after 60 minutes of AprA or polyP treatment. x indicates one protein upregulated in AprA but downregulated in polyP, one protein downregulated in both, and 19 proteins upregulated in both. (**B**) Venn diagram of significantly regulated phosphoproteins whose phosphorylation changes significantly relative to their total protein abundance at 60 minutes. y indicates one phosphoprotein upregulated in polyP but downregulated in AprA, and six phosphoproteins downregulated in both. (**C**) Venn diagram of unique phosphosites significantly altered by AprA or polyP at 60 minutes. z indicates one site upregulated in AprA but downregulated in polyP, two sites upregulated in polyP but downregulated in AprA, and six phosphosites downregulated in both. Numbers in each diagram represent proteins, phosphoproteins, or phosphosites that are significantly upregulated (above dashed line) or downregulated (below dashed line).

Phosphoproteomics profiling of the AprA-treated samples showed that among 602 quantified phosphoproteins, at 60 minutes AprA increased levels of 24 phosphoproteins including one (RRM domain-containing protein) with more than 2-fold upregulation and decreased 83 phosphoproteins including 16 (**Figure 1C, Supplementary file S3, tabs 2-4**) downregulated by more than 2-fold. At 60 minutes, polyP increased levels of 31 phosphoproteins and decreased the level of 35 phosphoproteins including 2 (Rac guanine nucleotide exchange factor T and SEC7 domain-containing protein) downregulated by more than 2-fold (**Figure 1D** and **Supplementary file S4, tabs 2-4**). For 6 phosphoproteins, both AprA and polyP decreased levels (**Figure 2B, Supplementary file S7, tab 7)**. One uncharacterized phosphoprotein (Primary accession: Q54Y28; gene ID: DDB_G0278453), which remains unannotated even after AlphaFold and FoldSeek searches, exhibited polyP increasing and AprA decreasing levels (**Supplementary file S7, tab 7)**, and none showed an opposite regulation.

### GO term analysis reveals overlapping and distinct pathways regulated by AprA and polyP

To explore pathways regulated by AprA and polyP, we performed Gene Ontology (GO) and SMART domain enrichment analyses using all significantly upregulated and downregulated proteins and phosphoproteins at the 60-minute time point. All enrichment results are provided in **Supplementary files S5** (AprA) and **S6** (polyP), including top enriched pathways, GO terms for biological processes, molecular functions, and cellular components, as well as associated SMART domains. All accession numbers for significant proteins, including both characterized and uncharacterized entries, were submitted to ShinyGO for analysis. For proteins and phosphoproteins listed as “uncharacterized” in Dictybase or UniProt, AlphaFold and FoldSeek were used to infer structural or domain information, and these annotations are summarized in the corresponding supplementary tables.

For AprA-regulated proteins, GO enrichment analysis revealed a dominant signature associated with RNA metabolism and ribosome biogenesis (**Supplementary Figure S1A–C and supplementary file S5, tabs 3-5**). Enriched biological processes included rRNA processing, small subunit of the ribosome (SSU)-rRNA maturation, ribosomal small-subunit biogenesis, mitochondrial RNA processing, and nucleolar RNA metabolism. Consistent with these functions, cellular component terms were highly enriched for nucleolus-associated ribosome assembly structures, including the 90S preribosome (a large precursor ribonucleoprotein complex that undergoes sequential processing to produce the small and large ribosomal subunits), small-subunit processome (a multiprotein complex required for early steps of SSU biogenesis), preribosome (a peripheral assembly intermediate associated with ribosome maturation) and ribonucleoprotein complexes, along with the nucleolus and nuclear lumen. GO molecular function analysis identified a single significant category, RNA binding (**Supplementary Figure S1D and supplementary file S5, tab 6**), suggesting an association between AprA-induced proteomic changes and transcriptional and translational control mechanisms.

PolyP-regulated proteins displayed a broadly similar enrichment profile, reflecting a shared impact on RNA-associated processes, but also exhibited distinctive features (**Supplementary Figure S2A–D**). Top biological processes included RNA surveillance and polyadenylation-dependent RNA catabolic pathways, as well as rRNA maturation and endonucleolytic cleavage events. Cellular component enrichment again highlighted ribosome assembly sites such as the preribosome, small-subunit processome, and nucleolus, alongside nuclear and membrane-enclosed lumens. Molecular function terms extended beyond RNA binding to include adenylyltransferase activity, suggesting that polyP signaling may influence RNA turnover and modification in addition to canonical ribosome biogenesis.

GO term analysis for the 21 proteins whose levels were affected by both AprA and polyP at 60 minutes showed a strong representation of ribosome biogenesis and RNA processing pathways, including endonucleolytic cleavage of rRNA, SSU-rRNA maturation, and ribosomal small-subunit biogenesis (**Supplementary Figure S3A–B and supplementary file S7, tabs 3-4**). Cellular component categories included nucleolar and ribosomal complexes (**Supplementary Figure S3C and supplementary file S7, tab 5**), while SMART domain enrichment identified ribosomal protein S1-like RNA-binding and helicase-associated domains (**Supplementary Figure S3D and supplementary file S7, tab 6**). Together, these results indicate that although AprA and polyP have largely distinct proteomic signatures, they converge on a functionally coherent group of proteins linked to ribosome assembly and RNA processing.

For AprA, enrichment analysis of the 107 phosphoproteins with significantly altered levels at 60 minutes (this includes phosphoproteins where change in levels of the phosphoprotein was due to changes in the total level of the protein; **Supplementary Figure S4A and supplementary file S5, tab 8**) highlighted categories related to RNA metabolism, cytoskeletal regulation, and small-GTPase signaling. GO molecular function analysis showed enrichment of guanyl-nucleotide exchange factor activity, mRNA binding, enzyme regulator activity and actin binding, pointing to actin remodeling and transcript control (**Supplementary Figure S4B and supplementary file S5, tab 9**). GO biological process enrichment analysis (**Supplementary Figure S4C and supplementary file S5, tab 10**) highlighted RNA decapping and methylguanosine-cap decapping, processes that remove the 5′ cap of mRNAs and thereby regulate transcript stability and turnover (Mugridge et al., 2018), suggesting that AprA can affect mRNA stability and processing. SMART domain analysis (**Supplementary Figure S4D and supplementary file S5, tab 11**) indicated enrichment of several domains such as Sec7 (ArfGEF) domains, pleckstrin homology (PH) domains, calponin homology domains and zinc fingers, indicating proteins that link membrane phosphoinositides to actin filaments (Korenbaum & Rivero, 2002; Lemmon, 2008). These patterns suggest that AprA may act through a combined regulation of RNA stability, protein degradation, and small GTPase signaling to promote chemorepulsion.

For polyP, enrichment analysis of the 66 phosphoproteins with significantly altered levels at 60 minutes (**Supplementary Figure S5A and supplementary file S6, tab 8**) included regulators of intracellular trafficking and GTPase signaling. GO molecular function terms (**Supplementary Figure S5B and supplementary file S6, tab 9**) showed strong enrichment for guanyl-nucleotide exchange factor activity, GTPase regulator activity and phosphatidylinositol binding. GO biological process enrichment (**Supplementary Figure S5C and supplementary file S6, tab 10**) indicated regulation of Arf protein signal transduction. SMART domain analysis (**Supplementary Figure S5D and supplementary file S6, tab 11**) identified enrichment of Sec7 domains (catalytic cores of Arf GEFs), zinc finger and DEAD-like helicase superfamily domains, underscoring involvement of Arf and Rho family GTPases in polyP signaling responses (Sztul et al., 2019). These data suggest that polyP affects levels of phosphoproteins controlling intracellular trafficking and small GTPase-dependent signaling (Ridley, 2006; Sztul et al., 2019).

We next examined phosphoproteins whose phosphorylation at 60 minutes changed significantly relative to the change in levels of the protein. For AprA (72 proteins; **Figure 2B**, **Figure 3A–D, and supplementary file S5, tabs 12-16**), GO terms highlighted regulation of Arf protein signal transduction, RNA binding, GTPase regulator activity, enzyme activator activity, and cellular components enriched at the cell cortex, cortical cytoskeleton and leading edge, consistent with phospho-regulation of actin-rich regions required for chemorepulsion. For polyP (25 proteins; **Figure 4A–D, and supplementary file S6, tabs 12-16**), enrichment included regulation of small GTPase-mediated signal transduction, mRNA cis-splicing via the spliceosome, phosphatidylinositol binding, guanyl-nucleotide exchange factor activity and snRNP nuclear complexes, suggesting links between polyP signaling, signal transduction, and RNA processing.

**Figure 3.**
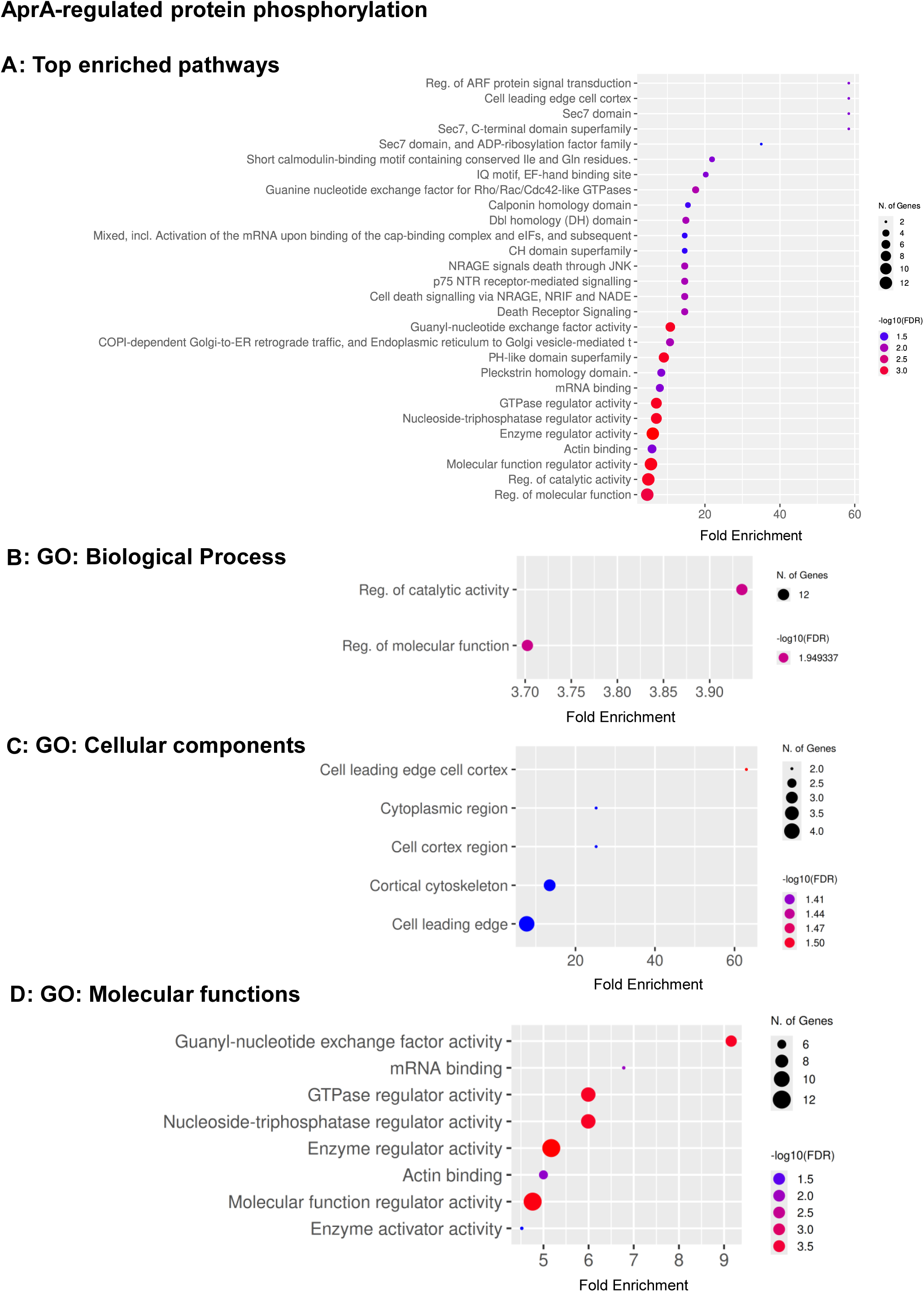
Functional enrichment of proteins whose phosphorylation is regulated by AprA. (**A**) Top enriched pathways, (**B**) GO: Biological Process, (**C**) GO: Cellular Component, and (**D**) GO: Molecular Function. Bubbles show fold enrichment (x-axis); bubble size represents the number of genes; bubble color indicates statistical significance as –log10(FDR), ranging from blue (minimum) to red (maximum). FDR cutoff = 0.05, pathway size = 5–1000.

**Figure 4.**
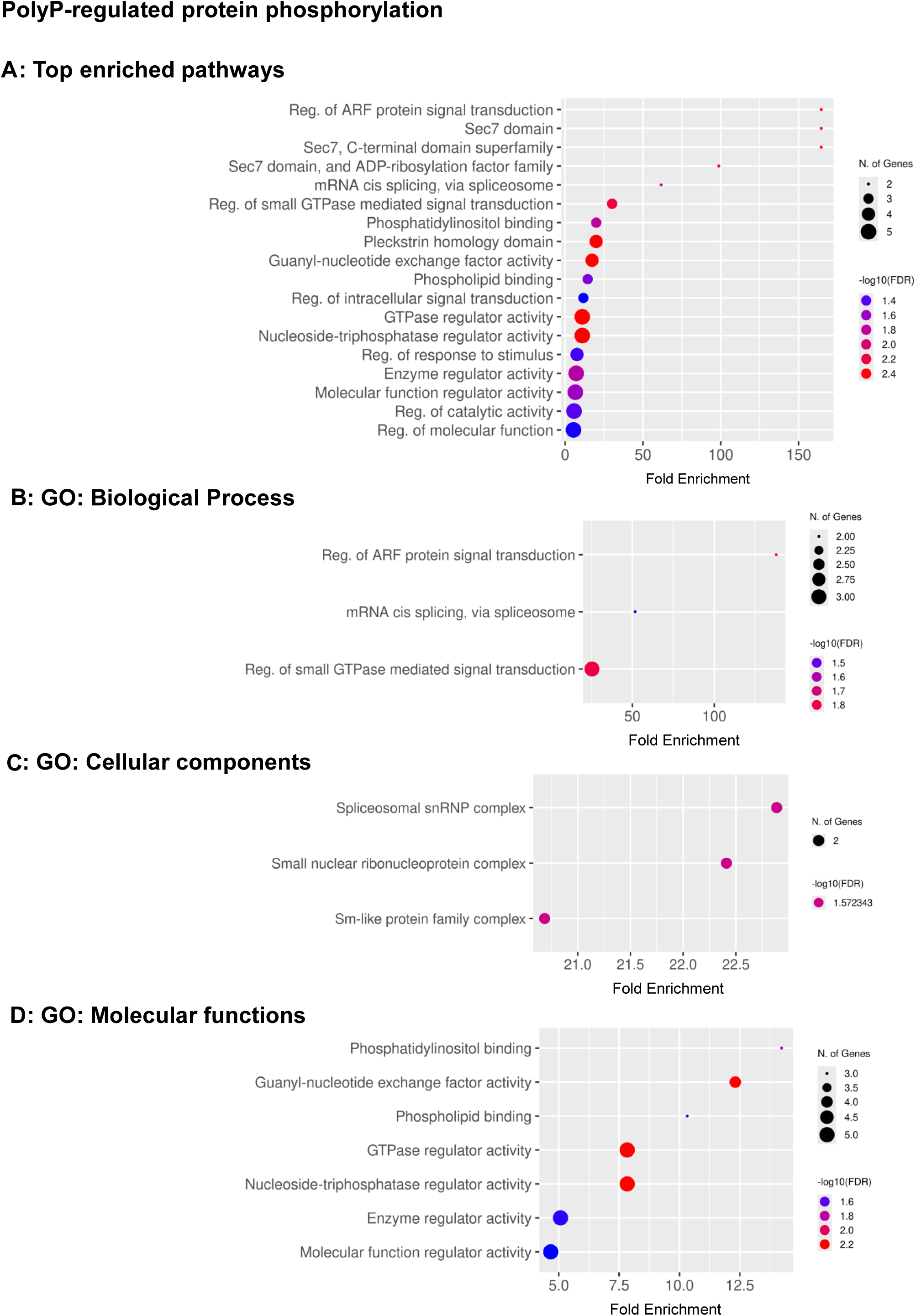
Functional enrichment of proteins whose phosphorylation is regulated by polyP. (**A**) Top enriched pathways, (**B**) GO: Biological Process, (**C**) GO: Cellular Component, and (**D**) GO: Molecular Function. Bubbles show fold enrichment (x-axis); bubble size represents the number of genes; bubble color indicates statistical significance as –log10(FDR), ranging from blue (minimum) to red (maximum). FDR cutoff = 0.05, pathway size = 5–1000.

The phosphorylation of 7 proteins was affected by both AprA and polyP (**Figure 2B and supplementary file S7, tab 7**). These included Sec7-domain (ArfGEF) proteins, the Ras-interacting protein RipA, the RacGEF GxcT, Rab-GAP–domain proteins, which are negative regulators of Rab GTPases that terminate vesicle trafficking cycles (Stenmark, 2009), and a Clu-domain–containing protein, which is typically involved in organizing multiprotein complexes important for signaling and organelle dynamics (Sen et al., 2015). Both AprA and polyP decreased the phosphorylation of six of these proteins, whereas the phosphorylation of one uncharacterized protein (Q54Y28; DDB_G0278453), which remains unannotated even after AlphaFold and FoldSeek searches, was decreased by AprA but increased by polyP. When individual phosphorylation sites were examined, AprA regulated 130 unique phosphorylation sites, whereas polyP affected 41 such sites (**Figure 2C and supplementary file S7, tab 8-10**). Among these, 9 sites were common for both treatments. Collectively, these overlapping proteins and their nine shared phosphorylation sites map to regulators of small GTPase signaling, membrane trafficking, and cytoskeletal remodeling, all core processes in directed cell movement (Hutagalung & Novick, 2011; Ridley, 2006; Sztul et al., 2019). Together, the results indicate potential points of signaling crosstalk and bifurcation between the AprA and polyP chemorepulsion pathways.

### Two proteins, GxcT and RipA, with repellent-decreased phosphorylation are necessary for chemorepulsion

To test the functional significance of phosphorylation changes identified in our phosphoproteomic analysis, we performed chemotaxis assays using strains with targeted knockouts of two candidate genes hypophosphorylated at 60 minutes after AprA or polyP exposure: *gxcT* (a Rac GEF; (Wang et al., 2013), and *ripA* (a Ras-interacting protein and TORC2 component; (Rosel et al., 2012). The primary metric was the forward migration index (FMI), which is the net displacement of cells along the gradient axis divided by the total path length (Foxman et al., 1999). Here, negative FMI indicates attraction toward the source, while positive FMI indicates repulsion. As previously observed (El-Sobky et al., 2025; Rijal et al., 2019) parental Ax2 (the parental strain for *gxcT^-^*) were repelled by AprA and polyP (**Figure 5**). The parental strain for *ripĀ* is KAx3 (Rosel et al., 2012), and we previously observed that KAx3 shows repulsion from AprA and polyP with no significant difference from Ax2 (El-Sobky et al., 2025; Rijal et al., 2019). In both AprA and polyP gradients, *gxcT^−^* and *ripĀ* cells showed no significant chemorepulsion (**Figure 5**). To determine if the lack of chemorepulsion of *gxcT^−^* and *ripĀ* cells was due to a general defect in cell motility, we measured migration speed and directionality (Euclidean distance divided by total distance) (**Supplementary Figure S6**). Although the mutants showed slight differences from Ax2 cells, they still showed motility, suggesting that the chemorepulsion defect in *gxcT^−^* and *ripĀ* cells is not due to a general impairment of motility.

**Figure 5.**
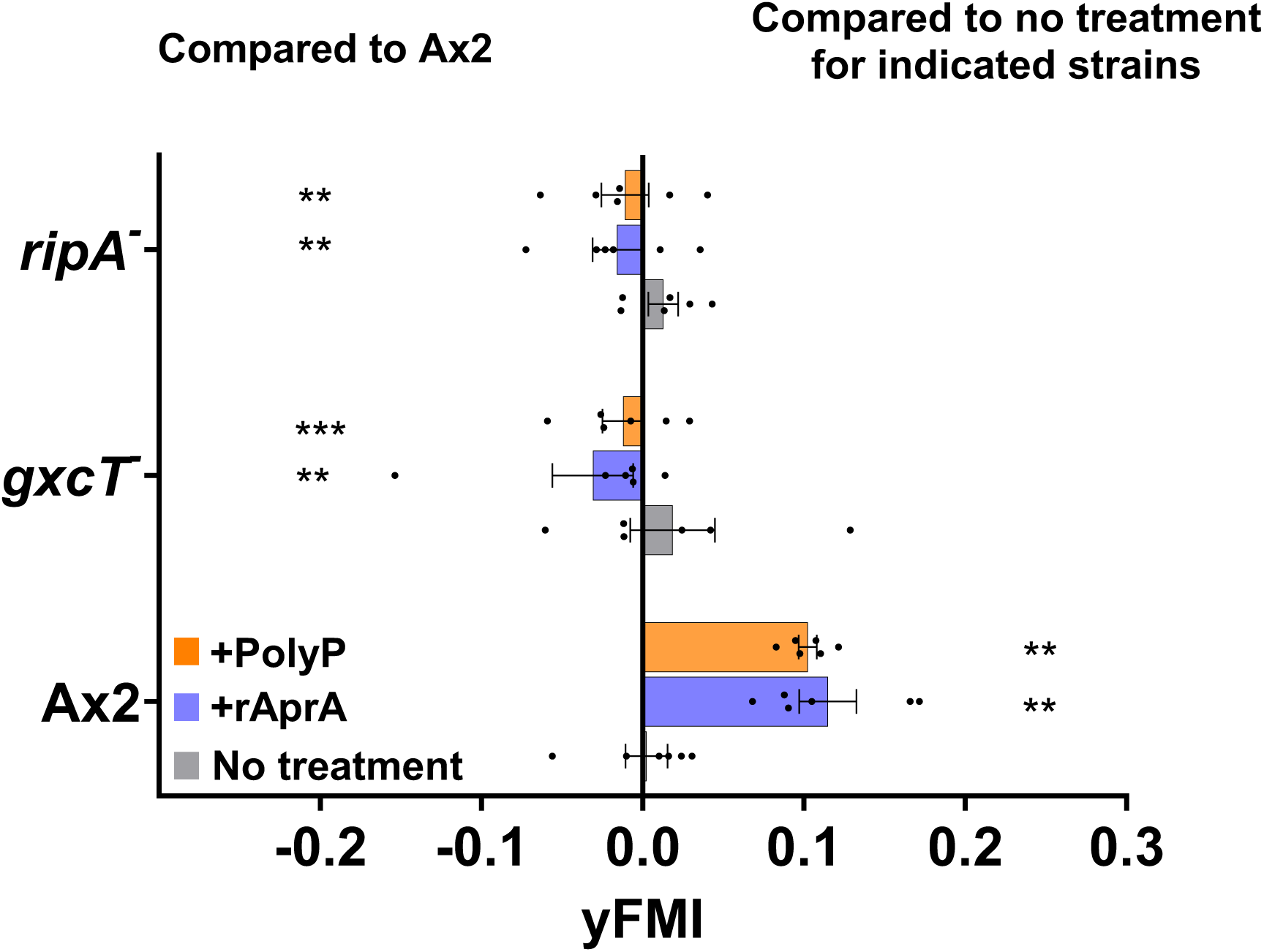
Chemotactic responses of phosphoprotein mutants in AprA and polyP gradients. Forward Migration Index (yFMI) along the gradient axis for wild-type Ax2 cells and phosphoprotein knockout strains *gxcT^−^* and *ripĀ*. Cells were exposed to AprA (blue), polyP (orange), or HL5 media as no treatment (gray). Positive yFMI values indicate repulsion from the source, while negative values indicate attraction. Bars are mean ± SEM, n ≥ 3. Statistical comparisons were performed using two-way ANOVA with multiple comparisons, analyzed relative to Ax2 (left column) and to untreated controls within each strain (right column). ** p < 0.01; *** p < 0.001.

### GxcT and RipA do not mediate AprA and polyP proliferation inhibition

In addition to inducing repulsion, AprA and polyP inhibit *D. discoideum* cell proliferation. As previously observed (Bakthavatsalam et al., 2009; Suess & Gomer, 2016), 300 ng/ml AprA and 700 µg/ml polyP reduced proliferation of Ax2 cells (**Figure 6**). Both the mutant strains showed proliferation inhibition by AprA and polyP similar to that of Ax2 cells, indicating that GxcT and RipA are part of the chemorepulsion mechanism and do not significantly mediate proliferation inhibition.

**Figure 6.**
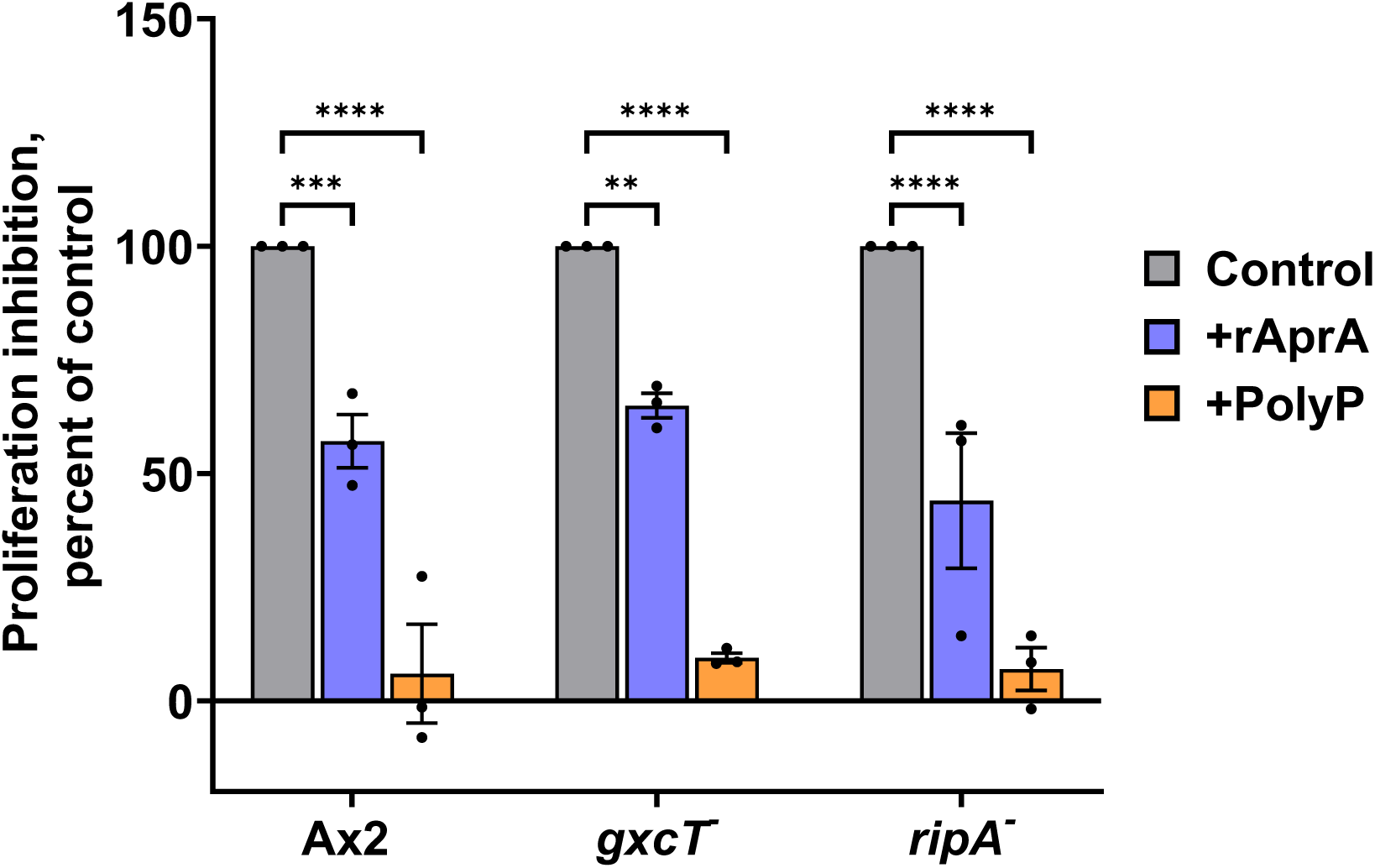
Proliferation inhibition analysis of wild-type and mutant *D. discoideum* strains in nutrient-rich media. Proliferation inhibition was quantified for the indicated strains after 24 hours in HL5 medium with AprA (blue), polyP (orange), or untreated control (gray). Values are the percentage of untreated controls. Bars represent mean ± SEM, n = 3. Statistical analysis was performed using two-way ANOVA with multiple comparisons. ** p < 0.01; *** p < 0.001; **** p < 0.0001.

## Discussion

Possibly as a way to deal with an extracellular environment where a signal might interact with the substrate and not diffuse properly, *D. discoideum* cells use two different chemorepellent signals to cause dispersal of cells from a growing colony. We previously found, by examining cell behavior and mutants lacking known signal transduction pathway components, that the two repellents use a combination of unique and overlapping pathway components to activate different chemorepulsion mechanisms. In this report, using an unbiased proteomics and phosphoproteomics approach, we find additional evidence supporting the differences in how AprA and polyP affect cells. In addition to the differences, there were some similarities, as the levels of a relatively small number of proteins, and the phosphorylation of a relatively small number of proteins, were regulated by both repellents. As with any proteomics or phosphoproteomics study, the quantitated proteins are only the most abundant proteins, and there are probably many more proteins whose levels and/or phosphorylation are regulated by AprA and/or polyP. Another important caveat in this study is that the cells were exposed to a constant level of the repellents, rather than a gradient, so there may be some different effects when cells are exposed to a spatial gradient rather than a constant level of the repellent. However, two of the proteins where both repellents decreased phosphorylation were necessary for repulsion by both repellents, indicating that for at least some proteins, a repellent-induced change in phosphorylation is associated with a functional role in repulsion.

At the edge of a colony, cells will be in a concentration of AprA or polyP that increases slowly as the colony size increases, but as these cells move away from the colony, the AprA or polyP concentration will decrease. Thus, unlike the pulsatile attractant cAMP, whose concentration at a given cell rises and falls within a few minutes (Gregor et al., 2010; Tomchik & Devreotes, 1981), the repellent concentrations change relatively slowly over time. After 60 minutes, we observed 72 proteins whose phosphorylation was affected by AprA, and 25 proteins whose phosphorylation was affected by polyP. Although this is considerably less than the almost 700 *D. discoideum* proteins whose phosphorylation was significantly altered by the chemoattractant cAMP (Nichols et al., 2019), it indicates that the chemorepellents affect a considerable number of proteins. For the proteomics, one unexpected result was that both repellents downregulated RNA metabolism proteins and upregulated ribosomal biogenesis proteins, suggesting that high levels of the two extracellular signals, indicating the presence of a large number of nearby cells, causes a cell to prepare to change its proteome from the proteome it had when it was a relatively isolated cell, perhaps derived from a dispersed spore. Both repellents affected, as expected, phosphorylation of signal transduction pathway components and unexpectedly affected phosphorylation of RNA processing proteins, albeit generally different sets of proteins. Combined with the effects on levels of ribosomal biogenesis proteins, the effects on phosphorylation of RNA processing proteins indicate that the two signals affect mRNA translation.

The phosphoproteomics profiles suggest that AprA and polyP influence distinct but partially overlapping signaling processes that could induce chemorepulsion. Both repellents reduced phosphorylation of proteins associated with Arf GTPases and Sec7-domain–containing guanine nucleotide exchange factors (ArfGEFs), which regulate vesicle trafficking, membrane remodeling, and actin cytoskeleton dynamics processes essential for directed cell migration (Donaldson & Jackson, 2011; Sztul et al., 2019). Despite this shared regulation, AprA inhibits pseudopod formation at the side of the cell closest to the source of AprA whereas polyP promotes pseudopod extension at the side furthest from the source of polyP (El-Sobky et al., 2025; Kirolos et al., 2021; Rijal et al., 2019). One apparent difference is that PolyP but not AprA affected phosphorylation of phosphatidylinositol binding proteins, which is consistent with the observation that polyP but not AprA requires phospholipase C and PI3 kinase for repulsion (El-Sobky et al., 2025).

Both *gxcT⁻* and *ripA⁻* cells failed to migrate away from AprA or polyP, indicating that these two proteins are required for chemorepulsion in response to both signals. GxcT encodes a Rho-family guanine-nucleotide exchange factor that stabilizes directional Ras activation and PIP₃ production during chemotaxis independently of the actin cytoskeleton (Wang et al., 2013). Its loss likely disrupts Rac-mediated cytoskeletal organization and polarity maintenance necessary for effective migration away from repellent cues. RipA (also known as Rip3/SIN1) acts as a Ras-interacting scaffold within the TORC2 complex, where it regulates F-actin dynamics and cell polarity (Khanna et al., 2016; Lee et al., 1999; Rosel et al., 2012). The loss of repulsion in *ripĀ* cells suggests defective TORC2-mediated cytoskeletal control downstream of Ras signaling. Together, these findings suggest that AprA and polyP converge on shared Ras- and Rac-GTPase modules through RipA- and GxcT-dependent regulation of TORC2 and cytoskeletal polarity, in addition to other signal transduction pathway components needed by both repellents (El-Sobky et al., 2025). Since the chemorepulsion of human neutrophils uses several of the signal transduction pathway components used by AprA (Consalvo et al., 2022), there is an intriguing possibility that human neutrophil chemorepulsion also involves some of the proteomics changes and phosphorylation changes identified in this report.

## Acknowledgments

We thank Dr. Alan Kimmel and Dr. Miho Iijima for gifts of cells. This work was supported by National Institutes of Health grant GM118355.

## Data Availability

The mass spectrometry proteomics and phosphoproteomics datasets, together with all supplementary files, are available on Figshare at Excel data files for AprA and polyP proteomics/phosphoproteomics

## Supplementary figure legends

**Supplementary Figure S1.**
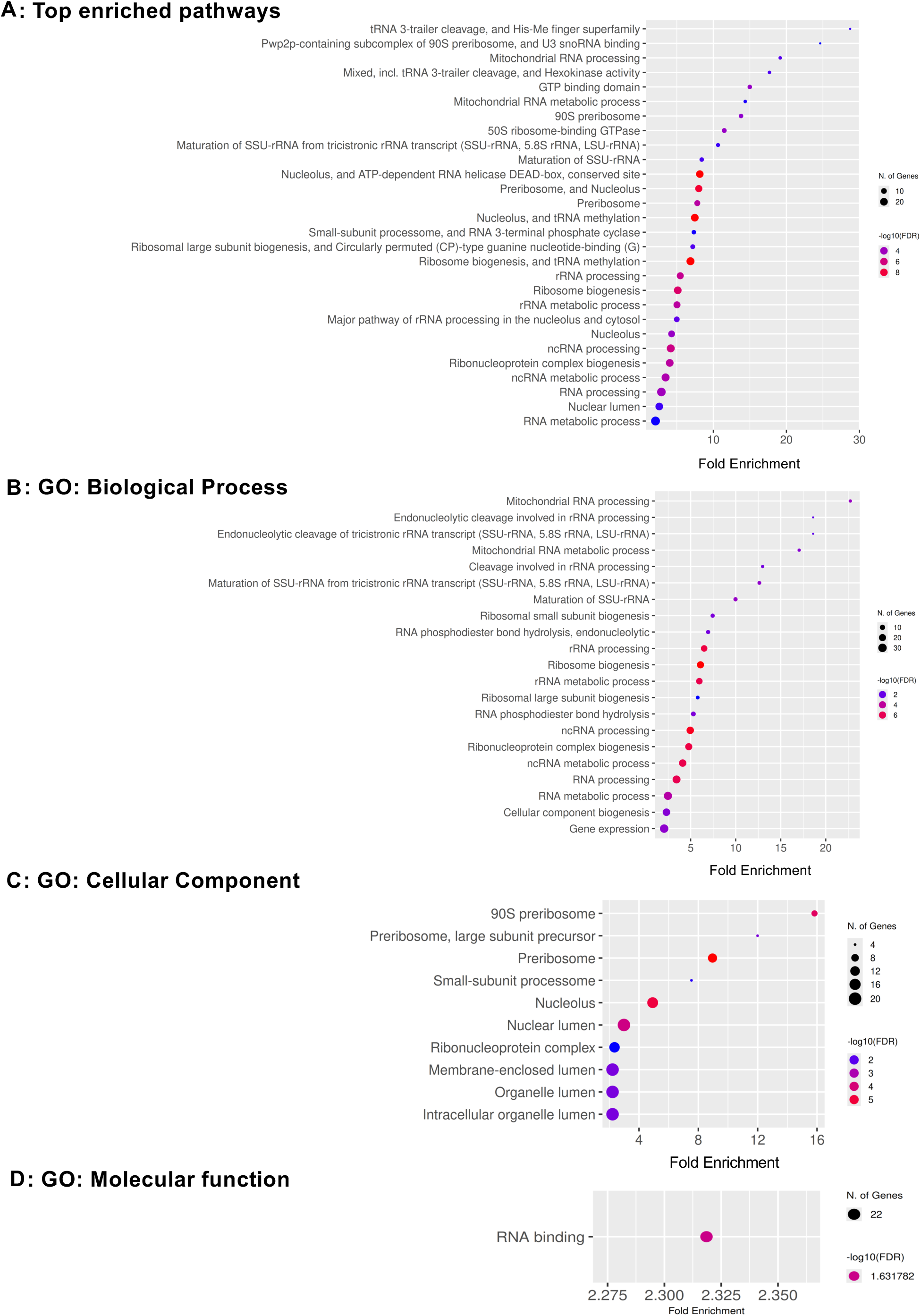
Functional enrichment of proteins whose levels are regulated byAprA. (**A**) Top enriched pathways, (**B**) GO: Biological Process, (**C**) GO: Cellular Component, and (**D**) GO: Molecular Function. Bubbles show fold enrichment (x-axis); bubble size represents the number of genes; bubble color indicates statistical significance as –log10(FDR), ranging from blue (minimum) to red (maximum). FDR cutoff = 0.05, pathway size = 5–1000.

**Supplementary Figure S2.**
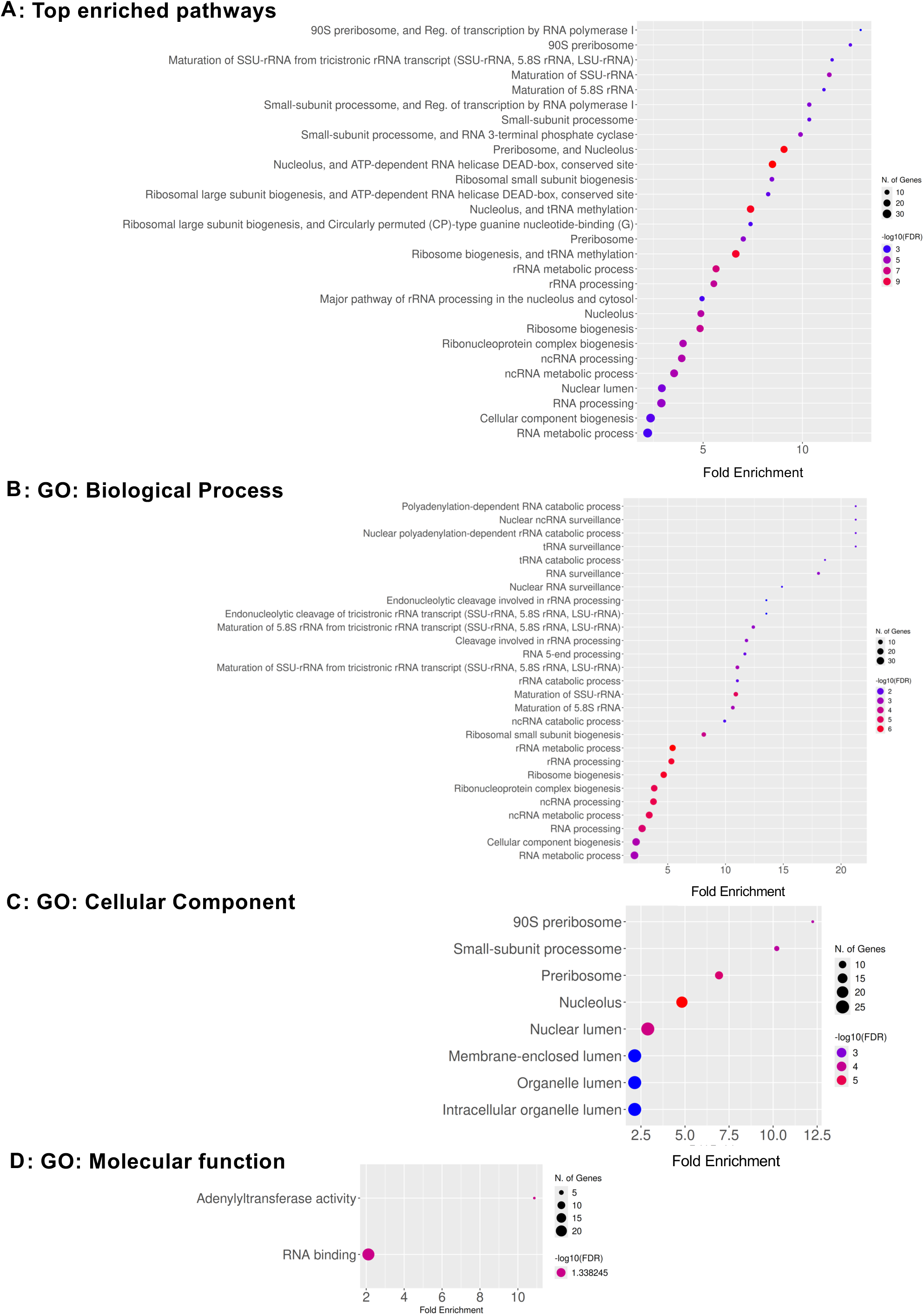
Functional enrichment of proteins whose levels are regulated by polyP. (**A**) Top enriched pathways, (**B**) GO: Biological Process, (**C**) GO: Cellular Component, and (**D**) GO: Molecular Function. Bubbles show fold enrichment (x-axis); bubble size represents the number of genes; bubble color indicates statistical significance as –log10(FDR), ranging from blue (minimum) to red (maximum). FDR cutoff = 0.05, pathway size = 5–1000.

**Supplementary Figure S3.**
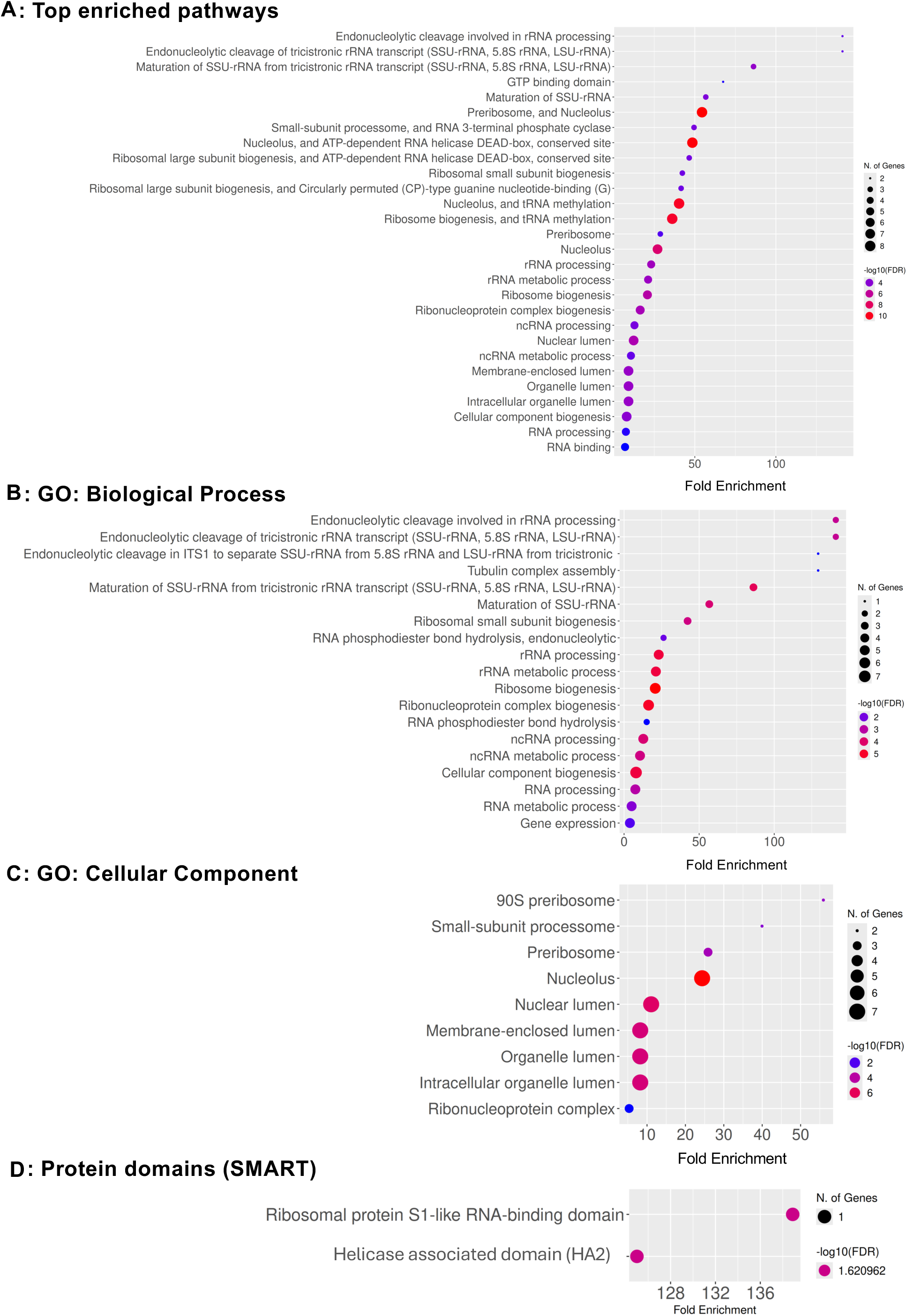
Functional enrichment of proteins whose levels are regulated by both AprA and polyP. (**A**) Top enriched pathways, (**B**) GO: Biological Process, (**C**) GO: Cellular Component, and (**D**) SMART protein domains. Bubbles show fold enrichment (x-axis); bubble size represents the number of genes; bubble color indicates statistical significance as –log10(FDR), ranging from blue (minimum) to red (maximum). FDR cutoff = 0.05, pathway size = 5–1000. No GO: Molecular Function was enriched for this group of proteins.

**Supplementary Figure S4.**
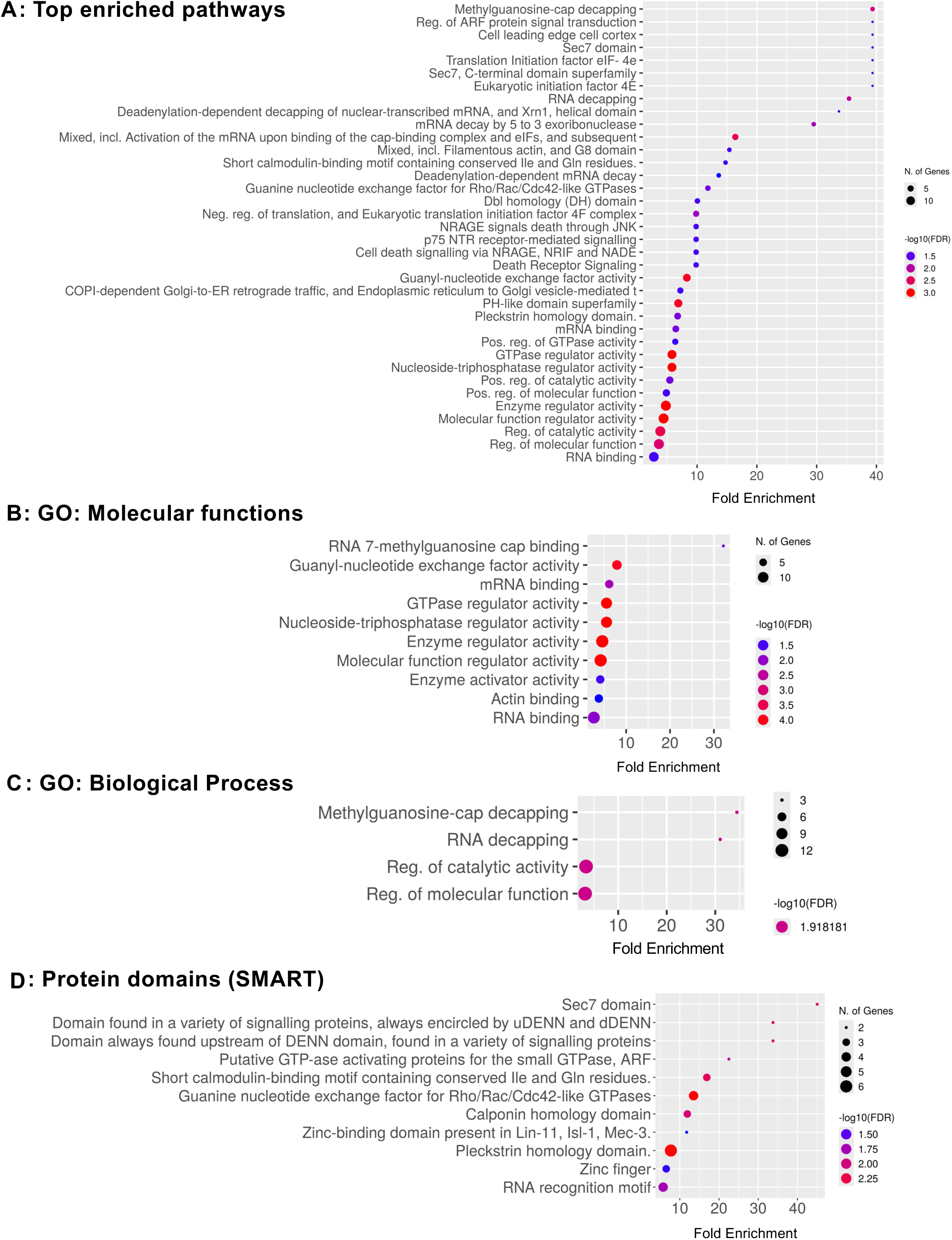
Functional enrichment of phosphoproteins whose levels are regulated by AprA. (**A**) Top enriched pathways, (**B**) GO: Molecular Function, (**C**) GO: Biological Process, and (**D**) SMART protein domains. Bubbles show fold enrichment (x-axis); bubble size represents the number of genes; bubble color indicates statistical significance as –log10(FDR), ranging from blue (minimum) to red (maximum). FDR cutoff = 0.05, pathway size = 5–1000.

**Supplementary Figure S5.**
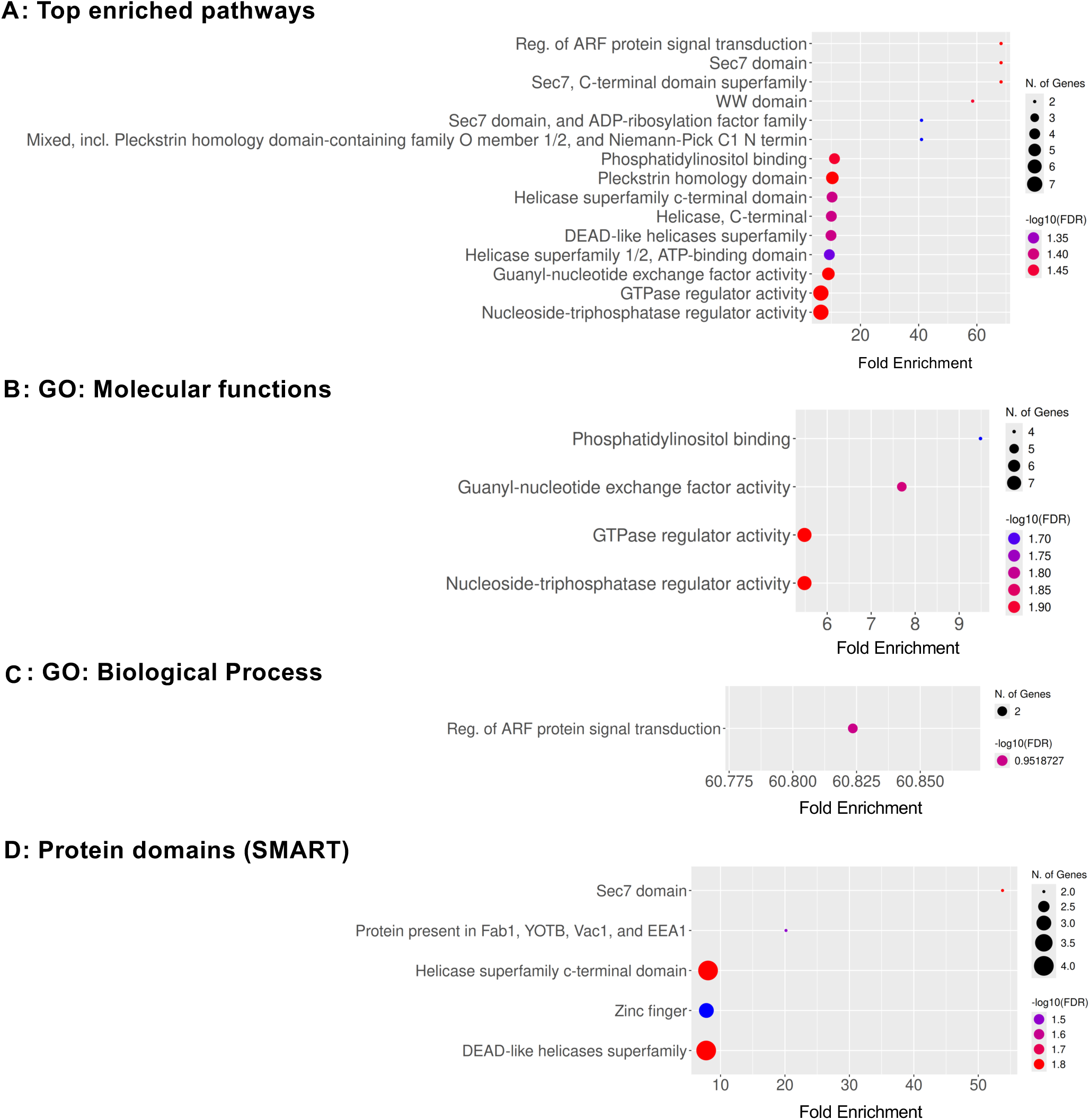
Functional enrichment of phosphoproteins whose levels are regulated by polyP. (**A**) Top enriched pathways, (**B**) GO: Molecular Function, (**C**) GO: Biological Process, and (**D**) SMART protein domains. Bubbles show fold enrichment (x-axis); bubble size represents the number of genes; bubble color indicates statistical significance as –log10(FDR), ranging from blue (minimum) to red (maximum). FDR cutoff = 0.05, pathway size = 5–1000.

**Supplementary Figure S6.**
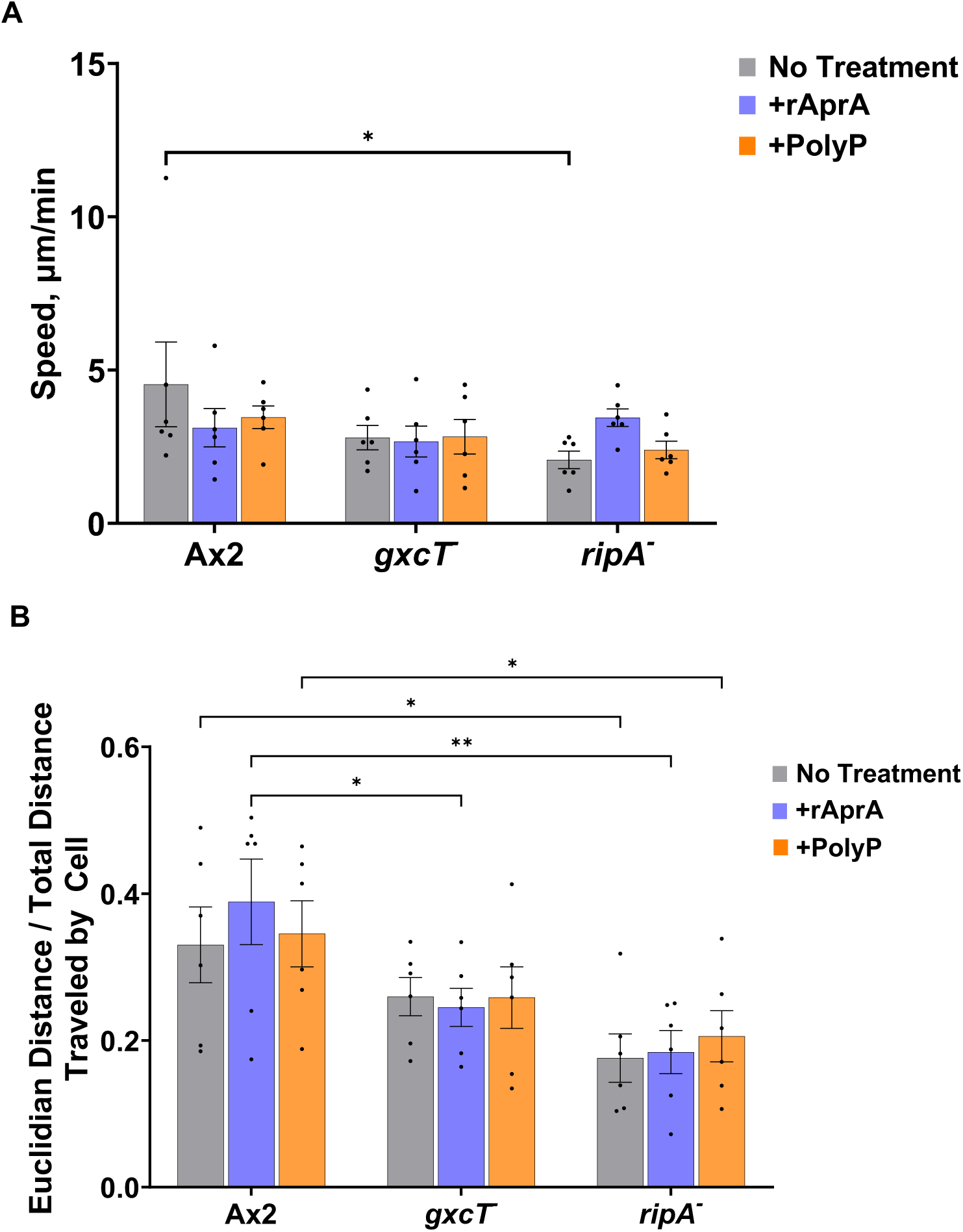
Chemotactic responses of phosphoprotein mutants in AprA and polyP gradients. (**A**) Average migration speed of the indicated strains. Bars are mean ± SEM, n ≥ 3. (**B**) Directional persistence of cell movement trajectories. Bars are mean ± SEM, n ≥ 3. Statistical comparisons were performed using two-way ANOVA with multiple comparisons. * p< 0.05, ** p < 0.01.

## Supplementary files

Supplementary File 1. AprA total proteins

Supplementary File 2. PolyP total proteins

Supplementary File 3. AprA phosphoproteins

Supplementary File 4. PolyP phosphoproteins

Supplementary File 5. Enrichment analysis of AprA-regulated proteins and phosphoproteins

Supplementary File 6. Enrichment analysis of PolyP-regulated proteins and phosphoproteins

Supplementary File 7. Proteins, phosphoproteins, and phosphosites regulated by both AprA and PolyP

